# Measuring chondrocyte viability of articular cartilage based on label-free two-photon microscopy and deep learning image analysis

**DOI:** 10.1101/2023.02.13.527931

**Authors:** Hongming Fan, Pei Xu, Xun Chen, Yang Li, Jennifer Hsu, Michael Le, Zhao Zhang, Emily Ye, Bruce Gao, Tong Ye

## Abstract

**Objective:** Chondrocyte viability (CV) is an important indicator of articular cartilage health. Two-photon excitation autofluorescence (TPAF) and second harmonic generation (SHG) microscopy provide a label-free method for imaging chondrocytes. In this study, we propose an automated assessment of CV using deep learning cell segmentation and counting based on acquired TPAF/SHG images.

**Design:** Label-free TPAF/SHG images of cartilage samples from rats and porcine were acquired using both commercial and home-built two-photon microscopes, respectively. TPAF/SHG images were merged to form RGB color images with red, green, and blue channels assigned to TPAF (two channels) and SHG signals, respectively. To make the training datasets for the deep learning networks, individual chondrocyte areas on the RGB color images were manually circled and live or dead chondrocytes were validated by using Calcein-AM and Ethidium homodimer-1 dye labeling. We first built a chondrocyte viability network (MCV-Net) using the Mask R-CNN architecture, which could provide individual segmented cellular areas with live or dead status. Wiener deconvolution preprocessing was added before the input of MCV-Net to improve the accuracy of the CV analysis, forming the Wiener deconvolution CV network (wMCV-Net).

**Results:** Training (300 images) and test (120 images) datasets were built for rats and porcine cartilage respectively. Wiener deconvolution could improve the Peak Signal-to-Noise Ratio (PSNR) for 30-40%. We demonstrated that both MCV-Net and wMCV-Net significantly improved the accuracy of the CV measurement.

**Conclusion:** A custom desktop TPAF/SHG microscope was used in collaboration with deep learning algorithm wMCV-Net based label-free method to assess the CV and get 95% accuracy with both rats and porcine samples.

## 1. Introduction

Articular cartilage provides a smooth, lubricated surface to absorb impact and distribute loads during movement so that underlying bone is protected. This function is facilitated by a complex and well-organized extracellular matrix (ECM) maintained exclusively by chondrocytes ^1,2^. Though they occupy a few percent volume of cartilage, chondrocytes play a critical role in maintaining a balance between anabolism and catabolism of matrix constituents such as water, proteoglycans (PGs), and collagens. A break of this balance may lead to osteoarthritis (OA), the most common joint disease affecting an estimated 10% of men and 18% of women over 60 years of age worldwide^3^. Chondrocyte death and survival are believed to be closely linked to cartilage breakdown^4^. As such, chondrocyte viability (CV), namely the fraction of viable chondrocytes in a cartilage tissue, is an important indicator of cartilage health. On the other hand, the CV of osteochondral allografts at the time of implantation is found to affect the longterm allograft survival rate^5–7^. CV is one of essential measures to reflect the quality of allografts. Methods to assess CV largely rely on introduction of dyes that can label live or dead cells specifically^8–10^ and allow either manual or automated cell counting to obtain CV. However, dye-labelled cartilage samples cannot be reused in further studies or in transplantation treatment procedures due to the potential cytotoxicity of dyes.

Autofluorescence of intracellular fluorescent coenzymes, such as reduced forms of nicotinamide adenine dinucleotide (NADH) or nicotinamide adenine dinucleotide phosphate (NADPH) and oxidized flavoproteins (FPs), have been long used as a label-free means to study metabolic states of cells^11,12^. Previous studies found that intensities of NAD(P)H and FP emission underwent significant variation when cells were transitioning from viable to nonviable states^13–15^, suggesting a potential label-free assay to distinguish live or dead cells. Our studies with freshly excised cartilage tissue from rat tibias^16^ showed that live chondrocytes exhibited higher intensity of autofluorescence from NAD(P)H than dead chondrocytes did. We found that the ratio between NAD(P)H and NAD(P)H+FPs emission was a robust measure to differentiate live and dead chondrocytes. For a more accurate viability analysis, we introduced second harmonic generation (SHG) images to reduce the interference from the autofluorescence of collagen in ECM. Images used in our viability analysis were acquired by a two-photon laser scanning microscope, which allowed collecting two-photon excitation autofluorescence (TPAF) from NAD(P)H or FPs, and SHG from collagen on the same set up. TPAF and SHG are both nonlinear optical processes involving two near infrared photons for excitation; we can simply name the related imaging modalities as two-photon microscopy under the category of nonlinear optical microscopy. It has been widely demonstrated that two-photon microscopy is suitable for imaging intact tissues excised or in vivo due to its large imaging depth and high spatial resolution. Taken all together, we proposed a label-free CV assay using images acquired by a two-photon microscope. We demonstrated that the sensitivity and specificity both exceeded 90% in the validation studies using intact rat tibias or fresh cartilage punches from porcine tibias. Similar to the dye-labelling assay, our label-free viability assay needs to visually count live and dead cells separately based on color appearance of chondrocytes in acquired images. This process requires human participation, and the analysis throughput is modest. To improve the throughput, automated techniques appropriate for analyzing cell images for measuring CV are desired.

Like many other cell-based image analyses, the required automated techniques involve two main image processing tasks: segmenting individual cell areas and classifying them to multiple categories (e.g., live and dead cells). Segmentation and classification are both classical problems in computer vision. Application of machine learning, especially, deep learning, to the computer vision has drastically advanced the field in terms of accuracy, speed, and complexity of tasks. Deep learning has been demonstrated in the segmentation of cells or cell components and classification according to various purposes^17^, yielding improved efficiency and accuracy in comparison with conventional algorithms. Recently, we have successfully demonstrated a deep learning approach for measuring CV with our label-free imaging assay^18^. In our method, we first built a U-Net model to segment lacuna areas on SHG images to generate masks for selecting areas containing cells. We then applied those masks to TPAF images to separate individual cell clusters. Two independent convolutional neural networks (CNNs) were developed to identify the number of live or the total number of cells in each cluster, respectively. CV was determined by summing up all live and total cells while going over every cluster. The reason for using two classification networks was to improve the accuracy since perfect segmentation of individual cells was difficult to achieve. Although a high accuracy was achieved in the viability analysis, the proposed method had an obvious disadvantage: multiple separate networks were required, and each network needed its own ground truth, model training, and evaluation processes. The overall workload and computing cost of retraining was high. Fortunately, the recently development of neural networks such as Mask R-CNN^19^ can include both segmentation and classification in a single architecture. Mask R-CNN is developed based on region-based CNN (R-CNN)^20^ and Faster R-CNN^21^ and provides a multitasking architecture that can perform object detection, target instance segmentation, and target key-point detection. Mask R-CNN is fast because its semantic segmentation (for finding masks), classification and bounding box regression run in separate branches in parallel. Mask-R-CNN uses the ROI Align technique to solve the pixel deviation problem in ROI Pooling and Fully Convolutional Network (FCN)^22^ to classify each pixel to the background or object categories. The pixel-level high accuracy in segmentation and classification makes Mask R-CNN a potential powerful neural network for microscopic image processing, where both image contrast and number of pixels are limited.

In this article, we demonstrate a Mask R-CNN based CV network (MCV-Net) specifically designed for measuring CV using images acquired using label-free two-photon microscopy. Through rigorous testing on images acquired from rat and porcine cartilages, MCV-Net achieves better accuracy than previously developed networks. We will use “previous CV networks” or pCV-Nets to indicate the previously developed multi-network deep learning method for CV analysis. We also demonstrate that Wiener MCV-Net (wMCV-Net), which adds a Wiener deconvolution^23^ in pre-processing before the MCV-Net, improves the accuracy of the viability measurement further. Taken together, we believe that the label-free two-photon microscopy and deep learning automated assessment provides a reliable, efficient tool for measuring CV in a non-destructive fashion.

## 2. Materials and Methods

### 2.1. Sample preparation

Rat and porcine cartilage samples were used in this study. For rat samples, entire tibias were excised from Sprague-Dawley rats (n=15) in accordance with the Institutional Animal Care and Use Committee’s regulations. After removal of muscles, tibias were held by a customized clamp prepared and tibia plateaus were aligned and imaged with settings as previously described^16^. For porcine samples, hind knee joints from adult Yorkshire pigs were obtained from a local meat processing company. Muscles were stripped away from the joint, and the joint cavity was opened to reveal the articular cartilage surface. Cartilage samples were harvested using 5 mm (ID) sample corers (18035-05, Fine Science Tools) and were stored in Dulbecco’s phosphate-buffered saline (DPBS, Corning) at 25°C. Cartilage punches were divided into 4 groups: 1) the fresh sample group, which were imaged right after harvesting; 2) the 4°C group, in which samples were kept in Dulbecco’s phosphate-buffered saline (DPBS, Corning) at 4 °C in a refrigerator; 3) the cultured group, of which samples were cultured in mixed culture medium (DMEM w/ Sodium Pyruvate, Penicillin-Streptomycin, Non-essential amino acids, and Fetal Bovine Serum) at 37°C in a VWR CO2 incubator (10810-944, VWR Air Jacketed CO2 incubator) for 3 days; and 4) the frozen group, of which samples were frozen in Dulbecco’s phosphate-buffered saline (DPBS, Corning) solution at −20 °C. All four sample groups were put in Petri dishes or 3D-printed sample holders and submerged in DPBS for imaging.

### 2.2. Label-free CV assay using two-photon microscopy

As we demonstrated previously, the label-free CV assay used a two-photon microscope to acquire three channel images: two TPAF channels for NAD(P)H and FPs, respectively, and one SHG channel for collagen in cartilage tissues. For visual assessment, we merged three channel images to RGB-colored images by assigning red, green, and blue colors to NAD(P)H, FPs, and collagen channels. Bright, green chondrocytes were identified as live cells while dim, red ones were dead cells. The viability of an imaged area was determined by calculating the percentage of live cells over the total cell population.

In this article, we included images acquired by two sets of two-photon microscopy systems, which were used to image rat and porcine samples, respectively. The two-photon microscope used for imaging rat samples was a commercial multiphoton laser scanning microscope (FV1200 inverted, Olympus Corporation, Tokyo, Japan) equipped with an ultrafast Ti:Sapphire laser (MaiTai Deepsee, Newport, CA) and two GaAsP photomultiplier tubes (PMTs). This microscope could only acquire two channels of images simultaneously. We typically tuned the laser to 740 nm first to acquire two channels of TPAF images and then tuned the laser to 860 nm to acquire SHG images. For TPAF imaging, the violet (420-460 nm) and red (575-630 nm) channels simultaneously recorded the autofluorescence of NAD(P)H and FPs, respectively. A 30x, 1.05 N.A. silicone immersion objective lens (UPLSAPO 30x, Olympus) was used to acquire images with a size of 1024 x 1024 pixels and a field of view (FOV) of 423 μm x 423 μm. A stack of 50 slices was acquired to cover a thickness of 50 μm of the cartilage tissue. For SHG imaging, the SHG signal at 430 nm was captured using the violet channel. TPAF and SHG imaging stacks were combined to form three-channel stacks using ImageJ (FIJI)^24^ for viability analysis. An example of such a set of TPAF and SHG images is shown in Fig. 1(A)-(D).

**Fig. 1.**
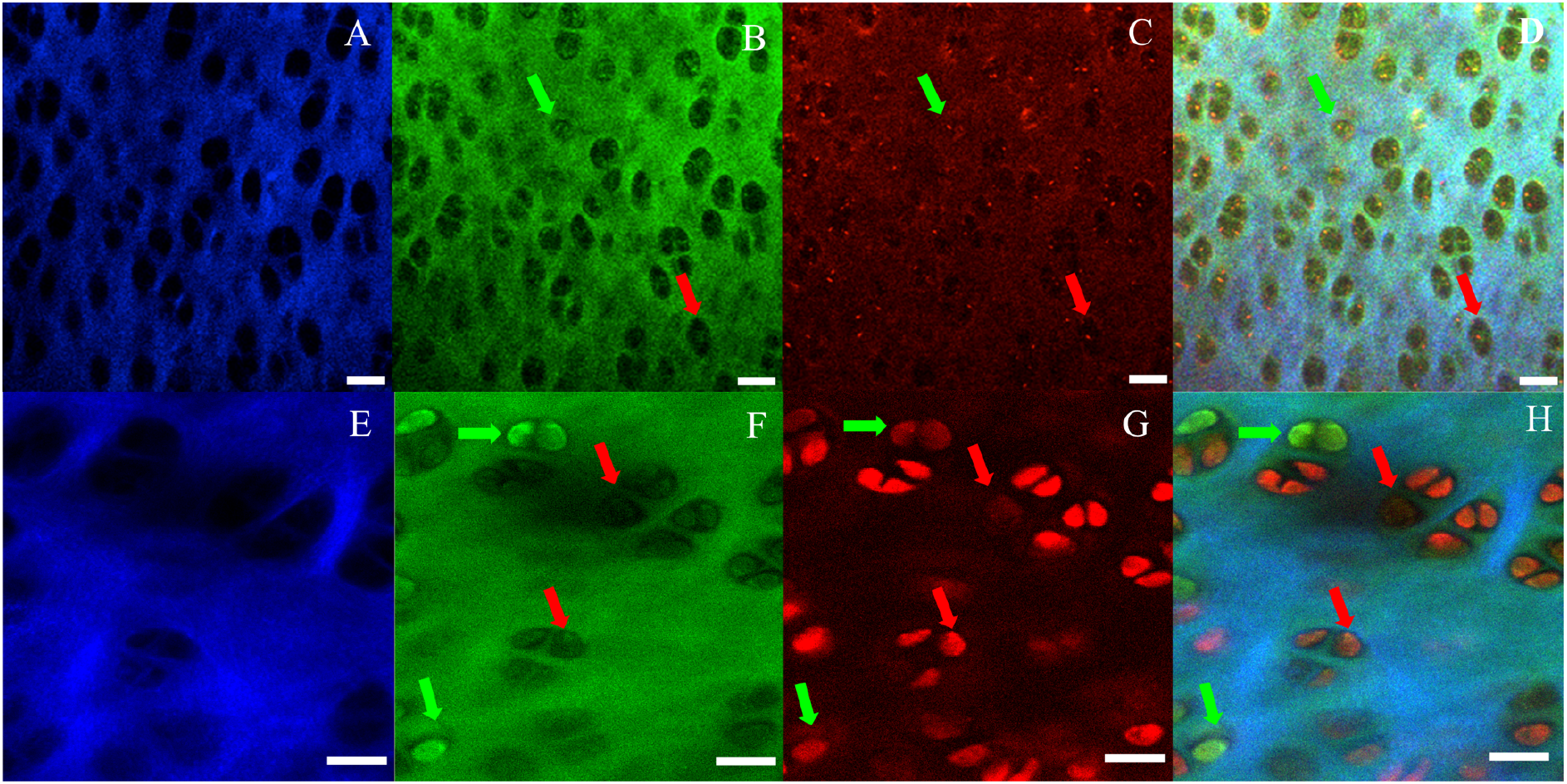
Typical pictures obtained by a commercial nonlinear microscope (FV1200) and home-build desktop microscope from rat cartilage tissue and pig cartilage tissue: (A)(a)SHG, (B)(b)Violet fluorescence,(C)(c) Red fluorescence, and (D)(d)merged RGB are examples of fluorescence. CV evaluation picture using (Red: red fluorescence; Green: violet fluorescence; Blue: SHG). Red arrows indicate dead cells, whereas green arrows indicate living cells. The images (A) (B) (C) and (D) in top line are from rat cartilage, and the images (a) (b) (c) and (d) in the bottom line are from pig cartilage. Scale bar:20μm.

The two-photon microscope used for imaging porcine samples was a home-built multi-channel two-photon microscope equipped with an ultrafast Ti:Sapphire laser (Chameleon Ultra II, Coherent Inc.) and two GaAsP PMTs (PMT2101, Thorlabs) and one Multialkali (PMTSS, Thorlabs). The system was designed with a focus on label-free imaging and the detail of the design was described previously elsewhere^25^. Three imaging channels on this microscope allowed us to acquire NAD(P)H, FPs, and SHG images pixel-by-pixel simultaneously. Similarly, the excitation laser was tuned to 740 nm while emission channels collected TPAF in the range of 421 – 463 nm (NAD[P]H) and 572 – 642 nm (FPs), and SHG in the range of 352–388 nm (Collagen). The imaging objective was 16x, NA 0.8 long working distance water dipping lens (CFI75 LWD 16X W, Nikon). Typically, at each location, an image stack containing more than 30 slices with a step size of 2 μm was acquired from the surface to deeper layers of a sample. Each image has a size of 512×512 pixels or a FOV of 150 μm x150 μm. TPAF and SHG imaging stacks were combined to form three-channel stacks using ImageJ (FIJI)^24^ for viability analysis. A set of TPAF and SHG images is shown in Fig. 1(E)-(H).

### 2.3. In situ dye staining assay and two-photon fluorescence imaging

The label-free CV assay was validated by using dye staining assays as described previously. In this study, the live/dead cell classification of rat samples was done with the validated label-free CV assay while the classification of porcine samples was done directly by dye-staining assay. Similar to the validation study, porcine cartilage samples were first imaged by collecting TPAF and SHG signals; then, leaving the samples untouched, two-photon excitation fluorescence imaging was performed on the same area after 30-minunte incubation with 2μM Calcein-AM and 8μM Ethidium homodimer (EthD-1). For imaging dye-staining samples, the laser was tuned to 800 nm and two-channel images were acquired with filters collecting emission in the range of 490 – 550 nm (Calcein) and 580-680 nm (EthD-1), respectively. Green and red color were assigned to Calcein and EthD-1 channels respectively for viability assessment. In this case, the green-colored cells were live while the red-colored ones were dead. However, in some cases, the dead chondrocytes are also shown in the green channel; as such, both channels need to be evaluated for the cell viability.

### 2.4. Training and test datasets

Similar to any deep learning methods, assembling annotated datasets including ground truth information for training is a critical and time-consuming part of model building. For Mask R-CNN based methods, the ground truth needs to include cell masks, mask locations, and cell status (live or dead). To determine the cell status, we performed imaging with live/dead dye assay described in the section of Materials and Methods. One thing we noticed that dead cells might appear in the green channel as well though, normally, they should appear only in the red channel on dye-staining images; however, those types of dead cells showed punctate patterns in the green channel. LabelMe^26^ was used to annotate TPAF/SHG images to include all needed ground truth information. The single cell area was manually circled with a polygon on a TPAF/SHG image. Live or dead status of a cell was determined by the appearance of that cell in the dye-staining image. Each annotated image was saved as JSON format. Two training data sets were built for rat and porcine samples respectively. Each training dataset consisted of 300, 8-bit RGB 3-channel, 512×512 TPAF/SHG images and their annotation files. Fig. 2 shows a typical TPAF/SHG image, its dye-staining image, and a screenshot of LabelMe during the annotation process.

**Fig. 2.**
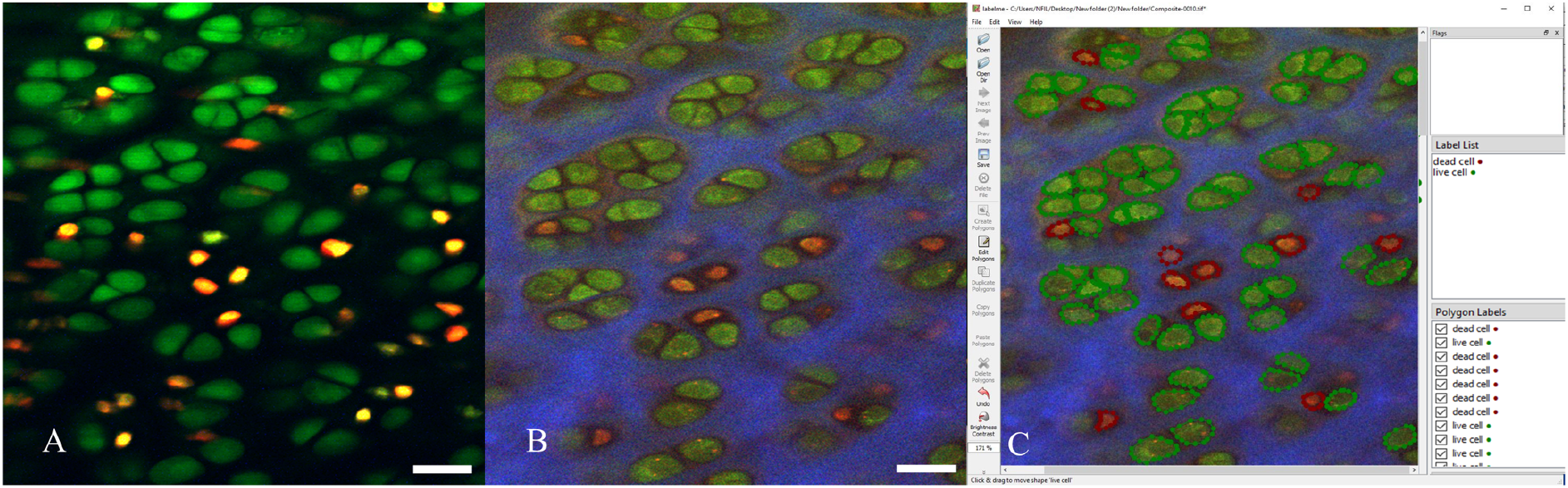
Training data made by using dye labeled images and nonlabeling images with python based software Labelme. (A) is the dye labeled images of pig cartilage, (B) is the Autofluorescence images of pig cartilage in the same location compare with dye labeled images. (C) is the window of software Labelme to label the cell with live and dead status based on dye labeled images.

For evaluation of the network performance, we built two test datasets for rat and porcine samples, respectively. The rat test set consisted of 120, 8-bit RGB 3-channel, 1024×1024 images from 50 individual image stacks (typically 30 images/stack) acquired from different tibia cartilage samples. The porcine test set consisted of 120, 8-bit RGB 3-channel, 512×512 images from 50 individual image stacks (typically 30-50 images/stack) acquired from different tibia cartilage samples. All images in the test dataset were annotated with LabelMe.

### 2.5. Deep-learning CV analysis using previously developed multiple networks (pCV-Nets)

For automated CV analysis, individual cells on TPAF images are segmented and classified as either live or dead based on appearance. The segmentation is difficult because both cell and the ECM regions give out comparable level of signals. Furthermore, chondrocytes are often nested in lacuna, in which a few cells stay close and touch each other, making it hard to separate individual cells. Considering these difficulties, the viability analysis strategy we previously proposed^18^ (pCV-Nets) consisted of two steps: 1) utilizing SHG images to generate masks for lacunae, and 2) using the generated masks to extract cell clusters on TPAF images for classification. We used a U-Net plus Watershed method to find the masks for lacunae. Our solution for extracting individual cells within cell clusters involve using two CNN networks to recognize the total number of cells and the number of live cells in each masked TPAF image. Each CNN network classified a cluster to one of five categories, which are identified by the number of all cells or live cells in the cluster with the minimum of 0, indicating no cells, and the maximum 4, indicating 4 and above number of cells. Three networks required three sets of training data sets. The U-Net model was built using Keras 2.1.6 library and TensorFlow 1.4.0 with over 4000 manually annotated inverted SHG images (256×256 pixels). The training procedure was completed on a customized GPU equipped with an Intel i9-7920x CPU and a graphics card (EVGA GeForce GTX 1080Ti FTW3).

Once the total number of cells and the number of live cells in each cluster were determined.

The CV (CV) was calculated by the following equation,

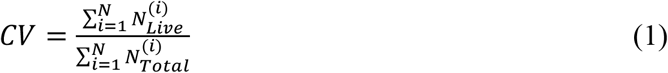

where *N_Live_* and *N_Total_* refer to the number of live cells and the total number of cells in the *i*-*th* cell cluster, respectively; N is the total number of cell clusters segmented in a measured (imaged) area.

### 2.6. Deep-learning viability analysis using a Mask R-CNN architecture

Mask R-CNN^19^ is the state-of-art method for instance segmentation, which labels each pixel to a known category of objects within an image. Mask R-CNN is suitable for counting objects that belong to different categories. In the case of CV measurement, Mask R-CNN outputs not only masks for each individual cell but also classification as dead or live for each mask. The output information is sufficient for calculating CV of an imaged region. Based on Mask R-CNN architecture, as shown in Fig. 3, Mask R-CNN is implemented with two consecutive stages. The first stage uses CNN^27^ to extract image features and the Region Proposal Network (RPN) to create a set of regions of interests (ROIs). The ROI pooling turns all these ROIs into fixed size and proposes the candidate object bounding boxes (BBoxes). Mask R-CNN uses ROI align (RoIAlign) to replace ROI pulling so that the exact spatial location can be preserved, and thus the accuracy of the mask is significantly improved. The second stage includes multiple branches running in parallel with high computing efficiency. The bounding box (BBox) regression branch determines if a BBox contains one of targeted objects according to the feature map with ROI Align mapping operation. The classification branch identifies which specific category the recognized object is in the BBox. The mask branch is a Fully Convolutional Network (FCN)^22^ that uses information generated from the BBox regression and classification branches to generate the mask for the region of objects in a pixel-to-pixel manner.

**Fig. 3.**
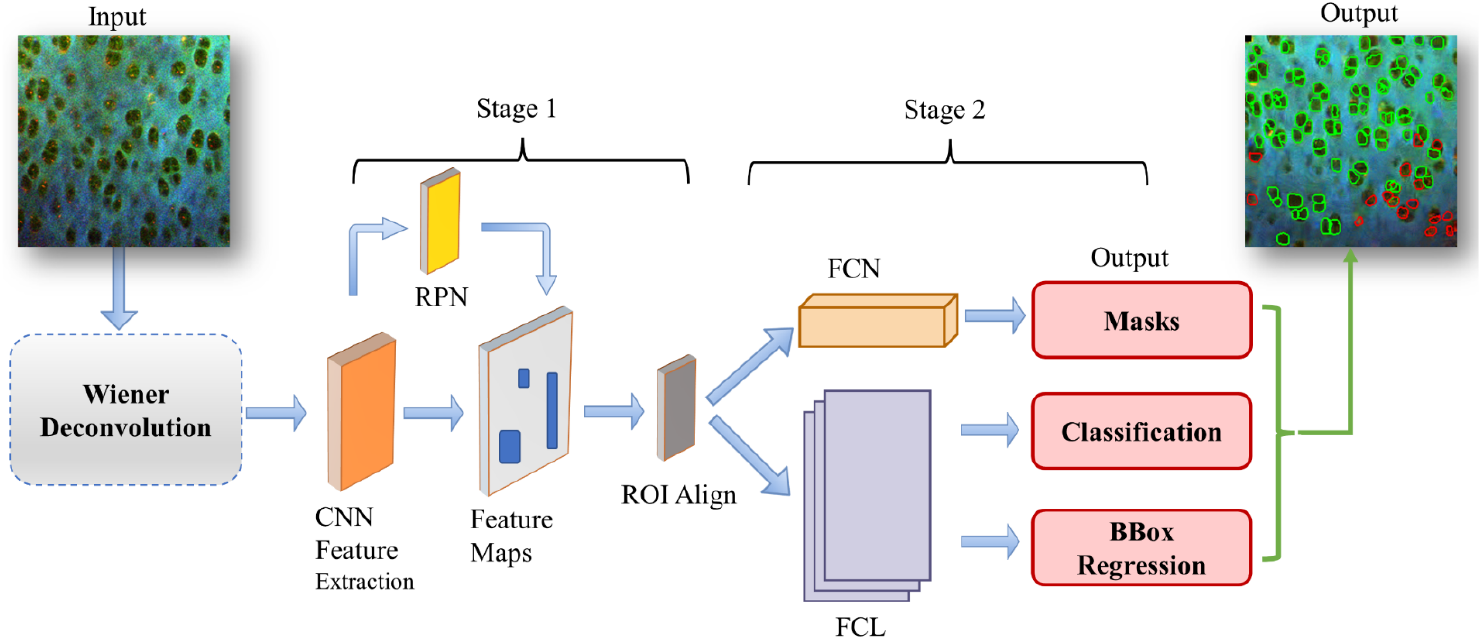
Wiener Deconvolution Mask R-CNN architectural diagrams for segmenting and categorizing chondrocytes. RPN is for region proposal network, FCL: Fully Connected Layers and ROI stands for region of interest.

The matplotlib 3.3, CUDA toolkit 10.2, Pytorch 1.8 and Torchvision 0.9 were used to implement the Mask-R-CNN model. The training set for the wMCV-Net algorithm included over 300 512 x 512 pixeled images with manually annotated chondrocytes. On a specialized computer with an Intel i9-7920x CPU and an EVGA GeForce GTX 1080Ti FTW3 graphics card, the training process took around 1 hour. The CV was calculated simply by dividing the number of live cells by the total number of cells.

### 2.7. Wiener deconvolution to improve the accuracy of viability analysis

To improve the accuracy of deep learning segmentation and classification, we introduced Wiener deconvolution as a pre-processing step to improve the image quality before the input of the deep learning network. Wiener deconvolution^23^ is one of classical, widely used image restoration algorithms that utilizes knowledge of the characteristics of the additive noise and the signal being recovered to reduce the impact of noise on deconvolution. It not only reduces image noise, but also eliminates image blur form image noise reduction, but also eliminate image blur caused by motion and other reasons. We used the following equation (see Supplementary Methods for more details) to implement Wiener deconvolution,

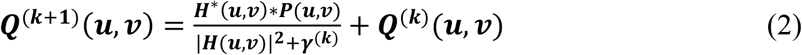

where ***H***(***u,v***) and ***H***\*(***u, v***) are the estimated transfer function and its complex conjugate of an imaging system; ***P***(***u,v***) is the power spectrum of an acquired image; ***Q***^(k)^(***u,v***) and *γ*^(*k*)^ are the ***k-th*** estimate of restored image and reciprocal of its signal to noise ratio (SNR). SNR and Peak Signal-to-Noise Ratio (PSNR) were used as measures to evaluate the improvement of an iteration cycle. Assume the image for restoration was an 8-bit one with a size of M×N pixels, the following equations were used to calculate SNR and PSNR^28^,

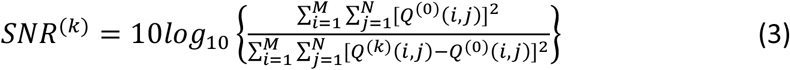

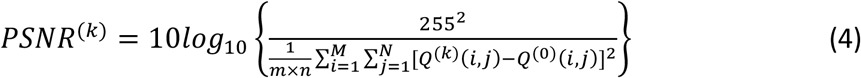

### 2.8. Performance evaluation of the CV measurement methods and statistical analysis

The performance evaluation of the CV measurement methods included two aspects: the performance of networks and the error of the CV measurement. For evaluating the performance of networks, we adapted a standard set of evaluation metrics commonly used in instance segmentation as follows.

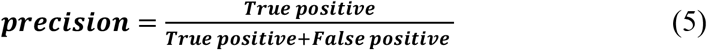

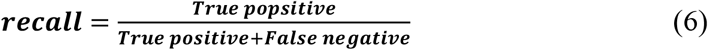

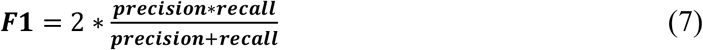

Note that, in the above equations, true-positive, false-positive, and false-negative indicate numbers of masks falling in each category and were determined differently when we compared the performance between previous multiple-network and Mask R-CNN based methods. For the multiple-network method (pCV-Nets), the evaluation was done by going over every cell cluster segmented by the U-Net. True positive was only assigned to those clusters that correctly predicted both the number of live cells and total number of cells according to the ground truth; otherwise, false positive was assigned. False negative was assigned to cell clusters where the algorithm failed to detect existing cells. For the Mask R-CNN based methods (MCV-Net and wMCV-Net), the evaluation was done by studying the intersection over union (IoU) of every mask of two classes (dead or live cells). IoU is a measure to quantify the intersection area between a predicted mask and a ground truth mask. The threshold of IoU was used to determine if a prediction was correct. A mask was defined as a true positive if its IoU is equal to or larger than 0.5 and otherwise a false positive. A false negative was an undetected mask. Precision reflects the fraction of correct predictions among all detected masks while recall (also known as sensitivity) reflects the fraction of correct predictions among all masks in the ground truth. By calculating the harmonic mean of a classifier’s precision and recall, the F1-score is a single value for reflecting the overall performance.

Additionally, we include the mean average precision (mAP)^29,30^ across classes to evaluate the performance of networks using the following equation:

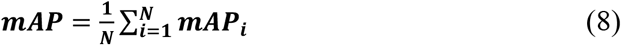

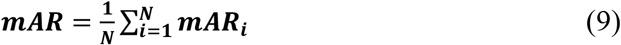

where *mAP_i_* and *mAR_j_* are the mean average precision (mAP) and the mean average recall^29,30^ (mAR) of the *ith* class, respectively, and N is the total number of classes. In the multiple network CV method, only one class was used to include the mask for cell clusters. In the Mask R-CNN based methods, two classes (live or dead) of masks were detected. For each class, AP is defined as the area under the precision-recall curve (PR curve) at a threshold value of IoU. While the IoU threshold is adjusted from 0.5 to 0.95 with an increment of 0.05, 10 APs, known as AP@[0.5:0.05:0.95], can be found and their average value is defined as *mAP* of a class (*see* https://cocodataset.org/#detection-eval). The final *mAP* is calculated by averaging through all classes.

Once the numbers of live and total cells are determined in each TPAF/SHG image, CV is calculated as a proportion of live cells ranging between from 0 to 1. The error of the measured CV was evaluated by the absolute error (AE) for each TPAF/SHG image and the mean absolute error (MAE) for an average over the number of test images. Additionally, the root mean square error (RMSE) is used to reflect the confidence of a measured CV. AE, MAE, and RMSE are defined as follows:

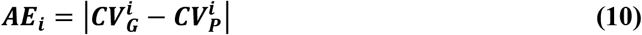

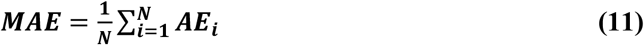

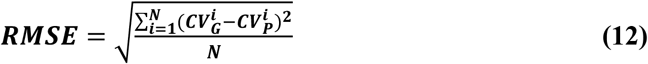

where 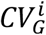 and 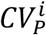 are the ground truth and predicted CV for the *i*-*th* image, respectively; N is the number of images in a test data set.

We conducted one-way analyses of variance (ANOVA)^31^ using Kruskal-Wallis and Friedman tests as well as t-tests, Mann-Whitney and Wilcoxon matched pairs tests. The processing group of the pig model (MCV-Net(wMCV-Net) and Mask-R-CNN) served as the independent variable for t-tests^32^, Mann-Whitney and Wilcoxon matched pairs tests. Processing group (U-Net, Mask-R-CNN, and MCV-Net(wMCV-Net)) of the rat model served as the independent variables in one-way analyses of variance (ANOVA) with Kruskal-Wallis and Friedman tests. For Sections 2.3–2.5, the dependent variables were accuracy and Pearson’s correlation coefficient, and P-values of less than 0.05 were regarded as statistically significant in all situations. GraphPad 9 (GraphPad Software, Inc.) was used to perform the statistical analysis.

## 3. Results

### Performance evaluation of Wiener deconvolution with TPAF/SHG images

To achieve the best possible accuracy for CV analysis, testing images (ones for evaluating networks) were pre-processed with Wiener convolution to reduce the noise level. To maintain the relative grey scale level between channels, Wiener convolution was simultaneously applied to all three channels of the RGB-colored images, which also meant SNR and PSNR of the colored images (not individual channels) were optimized through iterations. One essential factor to apply Wiener filter is the number of iterations to reach the optimal criterion measure for a specific application^23^. Using the test image sets, we studied the number of iterations required to reach the peak for images in the test datasets. As shown in Fig. 4, the average numbers of iterations to reach the peak between 45 and 50 for rat and porcine test images, respectively; the average PSNR improvement ratios are 40% for rat images and 30% for porcine images.

**Fig. 4.**
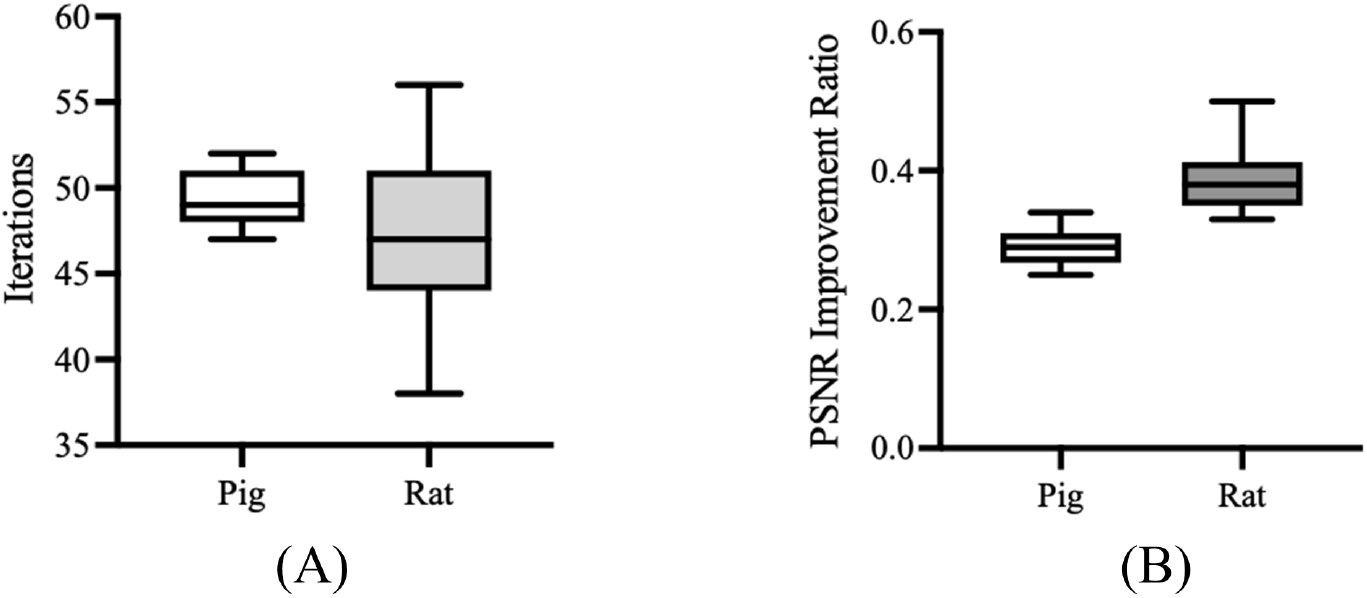
Wiener deconvolution processing result about the rat and porcine cartilage image iterations and PSNR (and SNR) improved ratio with statistic. (A) is the iterations statistic in rat and porcine cartilage images which Wiener deconvolution usually made (B) is the statistic about PSNR and SNR improved ratio in rat and porcine cartilage image with Wiener deconvolution processing.

The PSNR of rat images is improved more significantly than porcine images, which is perhaps due to a better contrast that the original porcine images have than the rat ones, consistence with the visual assessment. As shown in Fig. 5, after Wiener deconvolution process, the resulted image becomes smoother indicating reduced noise level.

**Fig.5.**
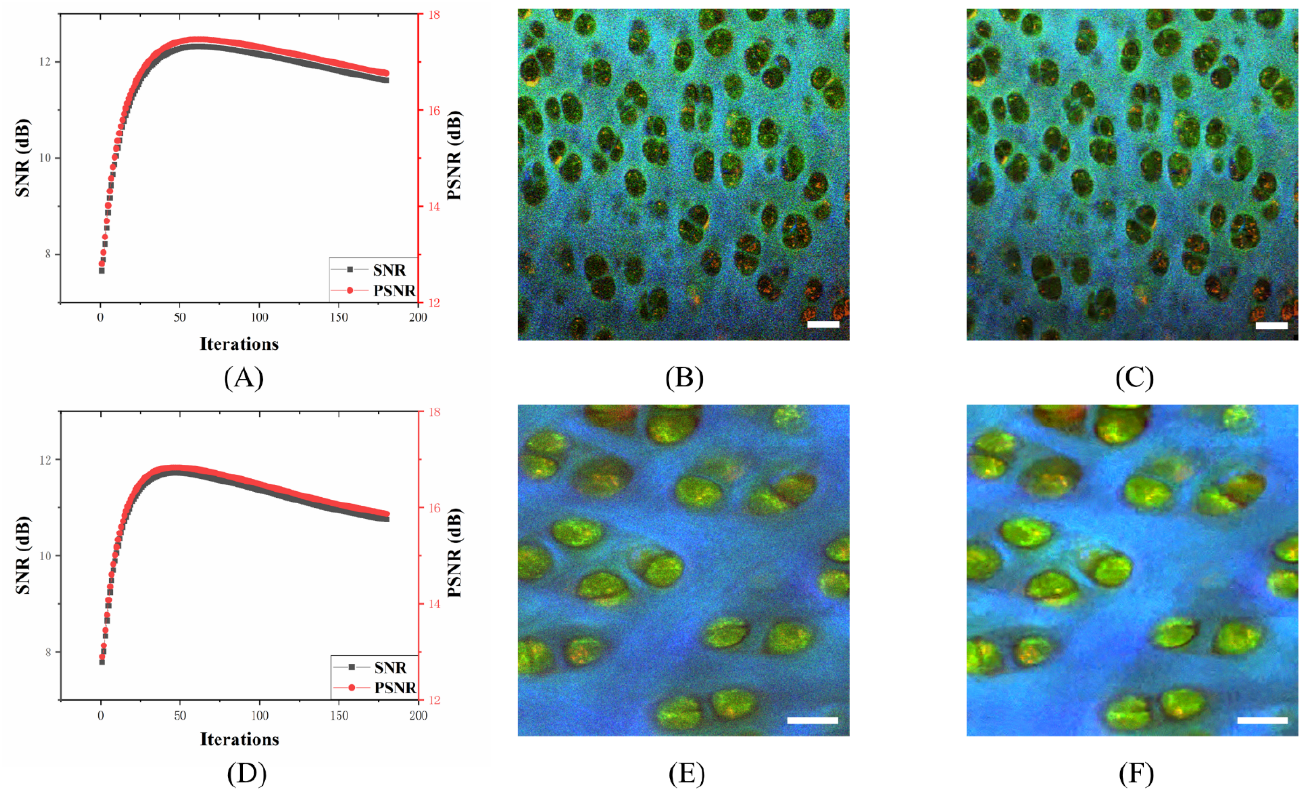
Image quality improvement with Wiener deconvolution processing. (A) is about the image quality relationship between iterations and PSNR (and SNR). (B) is the original raw rat cartilage image acquired by the commercial two-photon microscope FV-1200; (C) is the output result with the Wiener deconvolution processing; (D) is the discussion about how iterations and PSNR (and SNR) relate to image quality. (E) shows the initial raw image of porcine cartilage taken by the desktop home-build two-photon and SHG microscope; (F) shows the outcome of the Wiener deconvolution processing;

### Performance of instance segmentation at single cell level

Mask R-CNN based CV networks provide instance segmentation results in processing TPAF/SHG images. The instance segmentation for the CV analysis means that both the individual chondrocyte area (mask) and cell status (live or dead) are in the output of the network. In this study, we used test image datasets acquired from both rat and porcine cartilage samples to evaluate the segmentation performance. We also studied if the noise reduction using Wiener deconvolution could improve the segmentation performance. Representative segmentation results for rat and porcine images with their ground truth are shown in Fig. 6(A)-(F). On the ground truth images, dead and live cells are circled by red and green colored polygons respectively. On the output images of networks, red circles indicate the mark for the dead cells while green circles for live cells. Both MCV-Net and wMCV-Net predict masks and cell status accurately while wMCV-Net seems to perform a little better with less missed or wrongly identified cells and more accurate masks. We calculated mAP, mAR, and F1 score for each test image and performed statistical analysis over the outcome from 120 test images of each species. Note that F1 score^30^ was calculated using averaged mAP and mAR for each species. The outcome of the statistical analysis is summarized in Fig. 7. One-way analysis of variance (ANOVA) suggests that wMCV-Net shows a significantly better mAP, mAR and F1 score than MCV-Net in either species, which demonstrates Wiener deconvolution improves the segmentation performance significantly.

**Fig.6.**
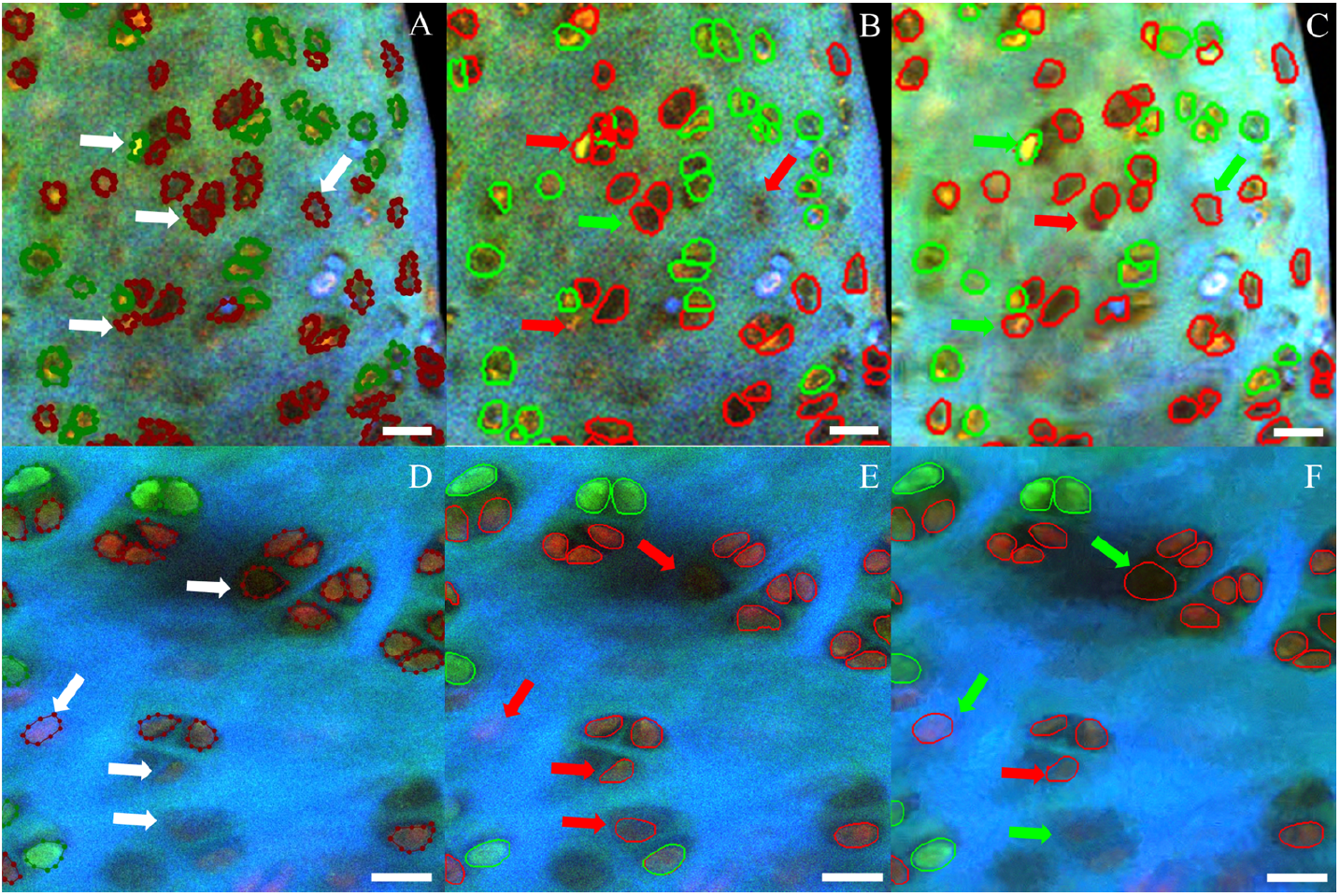
Performance of MCV-Net and wMCV-Net on rat and porcine cartilage images data. (A) is the ground truth of the rat cartilage image labeled by expert; White arrows stand for the standard of ROI segmentation and classification. (B) is result of rat cartilage image processed by MCV-Net; (C) is about segmentation classification result of rat cartilage image used by wMRC-Net. (D) shows the reference ROI segmentation and classification standard for the porcine cartilage image tagged with the corresponding dye labeled image. (E) shows the outcome of MCV-Net processing a porcine cartilage image; (F) shows the segmentation classification outcome of wMRC-Net processing the same porcine cartilage image. The red arrows indicate errors when compared to the standard ROI, which is shown by white arrows and green arrows represent the right result.

**Fig.7.**
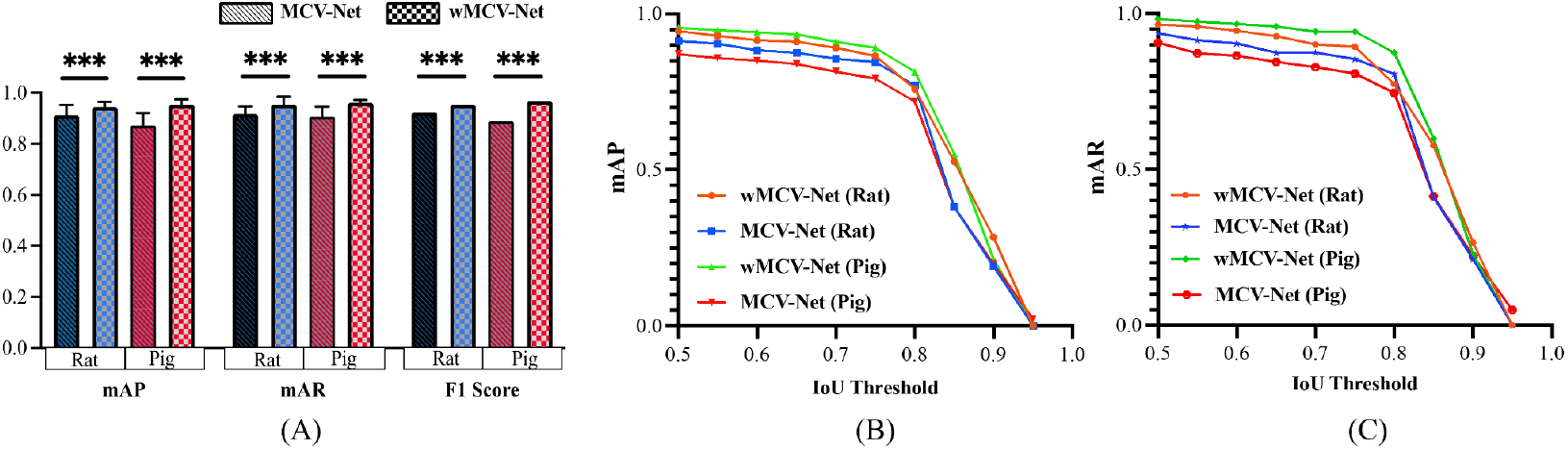
Comparison of segmentation accuracies achieved by MCV-Net and wMCV-Net on rat and porcine cartilage samples. On the basis of the percentage of matched and missing masks, the average accuracy score is calculated. (A) reported values to get a sense of the range of mAP, mAR and F1 scores with IoU threshold equal to 0.5. (B), (C) tested using specialized data (n = 240 test images, rat and porcine cartilage images) and quantified performance of the models MCV-Net and wMCV-Net. The IoU threshold measures the degree to which a predicted mask matches a ground-truth mask; a value of 1 denotes a pixel-perfect match, while a value of 0.5 denotes the proportion of successfully matched pixels to missed and false positive pixels.

The mAP, mAR and F1 score for either species exceeds 0.9 except for those for porcine test dataset using MCV-Net, which demonstrates Mask R-CNN based methods performs well in cell-based analysis. Fig. 7(B) and (C) shows mAP- and mAR-IoU curves respectively. Broader and taller curves can be seen for wMCV-Net in either species.

### Performance comparison of the networks for CV measurement

The goal of this study was to test if Mask R-CNN based CV networks (MCV-Net and wMCV-Net) could provide at least the same performance as the previous CV networks (pCV-Nets). respectively. Again, we used the test datasets from rats to compare the MAE to compare the accuracy of the CV measurement using three abovementioned networks. As shown in Fig. 8, the MAEs of pCV-Nets, MCV-Net and wMCV-Net are 0.14 ± 0.06, 0.08 ± 0.05, and 0.01 ± 0.02, suggesting that Mask R-CNN based networks are significantly better than pCV-Nets. The results demonstrate that Wiener deconvolution shows a significantly improvement of the accuracy in CV measurement; an accuracy of 0.95 is achieved.

**Fig.8.**
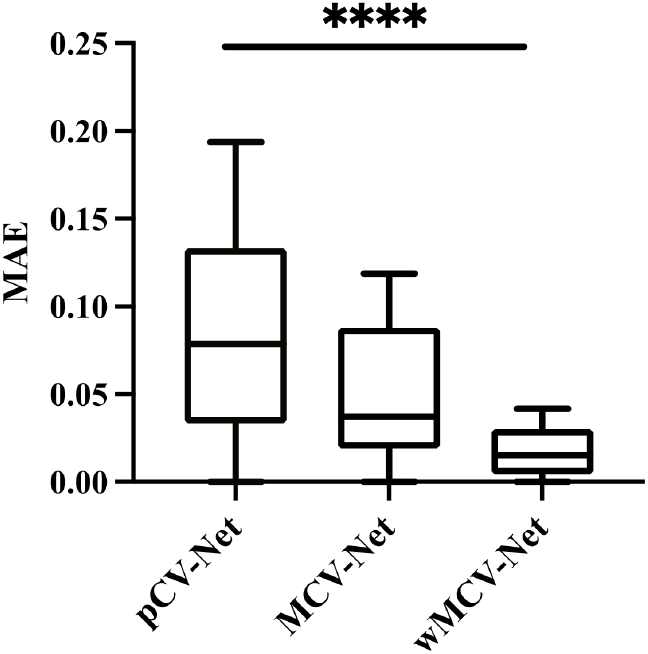
Root squared errors of viability examination employing cell number ratio-based deep learning techniques, integrating chondrocyte recognition techniques U-Net, MCV-Net and wMCV-Net with various animal species. At p <0.05, differences were considered as statistically significant. (****: p-value<0.0001)

## 4. Discussion

In this report, we demonstrate a novel approach for assessing chondrocyte viability using Mask R-CNN based networks for cell segmentation and classification. The new method has two significant improvements. Firstly, instead of using separate networks for segmentation and classification, MCV-Net is a single integrated architecture that can identify individual cells with their live or dead status. Rather than requiring multiple ground truth datasets using pCV-Net, MCV-Net only needs a single annotated dataset, making the training and viability analysis more efficient. Secondly, MCV-Net outputs masks for each individual cells while the previous method outputs masks for cell clusters. In pCV-Nets, the region of lacuna is segmented on SHG images and founded masks are used to extract cell clusters within each lacuna on TPAF images for the CV analysis. Although this strategy relaxes the strictness of segmentation of individual cells for maintaining high accuracy, the further single cell-based analysis is difficult. According to our two-photon images of articular cartilage, pericellular matrix (PCM) gives no or weak signals in either SHG or TPAF; SHG signals cannot exactly delineate the cell regions. MCV-Net marks the region of cells based on TPAF images and more precisely separates cell regions from the ECM than pMCV-Nets. Since the CV analysis only requires the cell counts in live and total cell populations, the accuracy of the cell region segmentation seems not that important; however, the argument is that missed cell regions may affect the feature recognition and thereby may affect the accuracy of cell classification as well. This is probably one of reasons that MCV-Net performs better than pMCV-Nets.

Due to the low quantum yields of NAD(P)H and FPs, TPAF images are often seen with low contrasts, which may affect accuracy of the CV analysis. To improve the image contrast, we studied one of commonly used noise reduction algorithms, Wiener deconvolution, to see if reduced noise level in input images can improve the accuracy of the viability analysis using MCV-Net. The results have shown that Wiener deconvolution improves PSNR 30%~40%. This contrast enhancement significantly reduces the error of the CV analysis (see Fig. 8 and Supplement Table S1). The number of iterations of Wiener deconvolution in our studies was set to 50 for all images to simplify the imaging processing procedure. However, as shown in Fig. 4, the number of iterations to reach the peak of PSNR varies for different images; it is possible to make an adaptive selection of the number of iterations according to certain criterions, which can improve the accuracy of the CV analysis further.

Since chondrocytes only appear in TPAF images, it is natural to think that the CV measurement should only need TPAF images and the SHG channel is not necessary. However, our studies shows that the SHG channel is necessary to improve the accuracy of identifying chondrocytes with its live or dead status. As shown in Table S2, including the SHG channel in the CV measurement significantly reduces the error when counting live, dead, and total cell numbers separately. The improvement is more prominent when using wMCV-Net. The contribution from the SHG channel was also found in our previous study, where populations of live and dead cells could be better separated according to the NAD(P)H and FPs ratios if the SHG channel was used to reduce the autofluorescence contribution from collagen in the ECM region^16^. The usefulness of the SHG channel in CV analysis is perhaps due to the overall improvement of the contrast between cells and the ECM area. As we can see in raw TPAF images (Fig. 1), signals from the ECM, especially in the NAD(P)H channel, is comparable with those from cells, making it difficult to decern cells in TPAF images. As seen in Fig. 1(D) and (H), the RGB image with the SHG channel assigned to blue color, improves the visibility of cells, and thereby improves the accuracy of the cell recognition. SHG images are highly recommended to be included in the CV analysis (Supplement Table S2 and Figure S1). If three channels are equipped on the microscope, the SHG imaging can be readily acquired simultaneously with TPAF channels without taking any extra time and effort.

MCV-Net and wMCV-Net are based on the Mask R-CNN architecture, which is one of the most widely used high performance algorithms for instance segmentation. MCV-Net or wMCV-Net segment pixels that belong to each individual chondrocyte. Although the CV analysis only requires cell classification and counting, MCV-Net and wMCV-Net also output masks for individual chondrocytes, which provides possibilities to expand the application beyond the CV analysis. For example, segmented cell areas can be used to determine cell sizes. Although we focus on the CV analysis using label-free imaging method in this study, the Mask R-CNN approach can be applied to cell-based applications using dye-labeling.

A few years ago, our team proposed a label-free (non-labeling) assay to identify live and dead cells for CV assessment in articular cartilage tissue. In this assay, intrinsic nonlinear optical signals such as TPAF and SHG are used to image cells and the ECM in high spatial resolution and in three dimensions. TPAF and SHG imaging together are usually called two-photon microscopy for both need two photons in excitation. The non-destructive and dye-free nature of our proposed assay allow us to measure CV on native tissues. Conducting validation studies using dyes on several thousands of cells from two species (rats and pigs), we have demonstrated that both visual assessment and quantitative analysis based on our label-free assay are as reliable as dye-based assays. Since the samples are reusable after imaging, the proposed assay may find applications in tissue preservation and quality control of allografts. In those types of applications, high throughput in the measurement is essential. Due to the relative low image contrasts, conventional image processing algorithms did not work well. Deep learning seems a promising solution for improvement of the throughput. As we demonstrated in this report, the Mask R-CNN based architecture, MCV-Net or wMCV-Net ultimately solved the throughput problem. Plus the customized two-photon microscopic imaging instrument and acquisition protocol (see Supplementary Data for more details), we believe that we present a complete package for measuring CV of articular cartilage tissues without dye labeling.

## 5. Conclusion

We have shown that live or dead chondrocytes of articular cartilage can be identified on label-free images acquired by a TPAF/SHG microscope. For measuring the chondrocyte viability, we have developed a Mask R-CNN based method, called MCV-Net, to segment individual cells and classify their live or dead status based on acquired label-free images. We further demonstrate that adding denoise preprocessing such as Wiener deconvolution on MCV-Net, can improve the accuracy of the CV measurement. Ultimately, wMCV-Net (Wiener deconvolution MCV-Net) can reach 95% accuracy in the CV measurement. We have demonstrated a complete CV measurement package, including a label-free imaging method and deep learning based automated cell recognition and counting.

## Author contributions

HF built the desktop homebuilt two-photon microscope, acquired all porcine images, and analyzed data. HF and PX wrote codes for implementing Mask R-CNN based algorithms for CV analysis. XC and JH coded the U-Net based multiple networks for CV analysis. YL acquired all rat images. ML and ZZ helped with image annotation and data processing. EY, BG provided clinical data interpretation. HF and TY conceived the original idea and wrote the manuscript. TY supervised the project.

## Supplementary data

See the attached document.

## Supplementary Materials

### Supplementary Methods

#### Deduction of 2D Wiener deconvolution

For the above case, the formula of Wiener’s deconvolution filter can be expressed as:

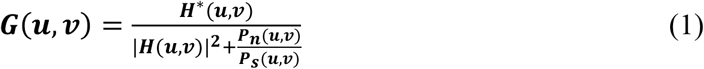

Where ***H***(***u, v***) is degenerate function, and ***H***\*(***u, v***) is complex conjugate of ***H***(***u, v***). ***P_n_***(***u, v***) stands for noise power spectrum, and ***P_s_***(***u, v***) represents the power spectrum of undegraded image. By defining the error function, we can get the following function:

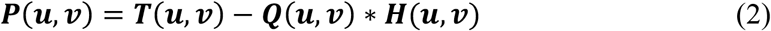

Where ***T***(***u, v***) is the degraded image. And Substituting function (2) into function (1) and replacing ***Q_old_***(***u, v***) and ***Q_new_***(***u,v***) with ***P***(***u,v***) and ***G***(***u,v***), we get:

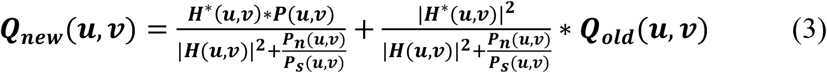

Eq. (3) is the iterative formula in the usual sense, which shows that one iteration is equivalent to superimposing an increment on top of the original, which is essentially an incremental Wiener deconvolution filter. Since ***P_n_***(***u, v***) is usually much smaller than ***P_s_***(***u, v***), the ratio can be expressed as a number γ less than 1, so that equation (3) can be approximated as follows:

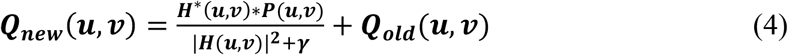

In addition, we can also get:

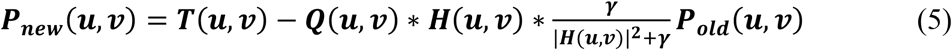

where ***P_old_***(***u, v***) denotes the error of the previous iteration. We can see that 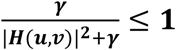, so ||***P_new_***(***u, v***)||^2^ ≤ || ***P_old_*** (***u, v***)||^2^. That is the iteration error steadily lowers as the number of iterations rises. However, excessive iterations can also result in picture quality loss owing to the effects of noise and the corresponding motion in the system.

### Supplemental Text/Video

**Fig. S1.**
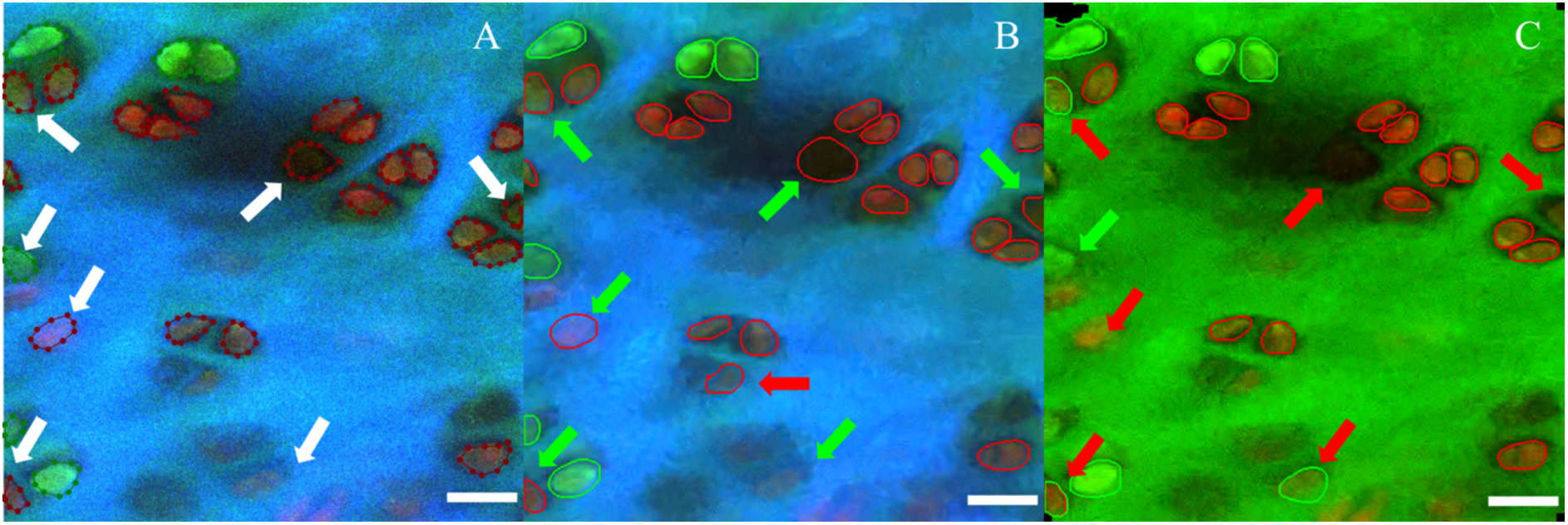
The performance about accuracy of identifying chondrocytes with its live or dead status in different channels. (A) is the ground truth of the porcine cartilage image labeled by expert; White arrows stand for the standard of ROI segmentation and classification. (B) is result of three channels (NADH, FAD and SHG) of the porcine cartilage image processed by wMCV-Net; (C) is about segmentation classification result of two channels (NADH and FAD) of the porcine cartilage image used by wMRC-Net. The red arrows indicate errors when compared to the standard ROI, which is shown by white arrows and green arrows represent the right result.

**Fig. S2.**
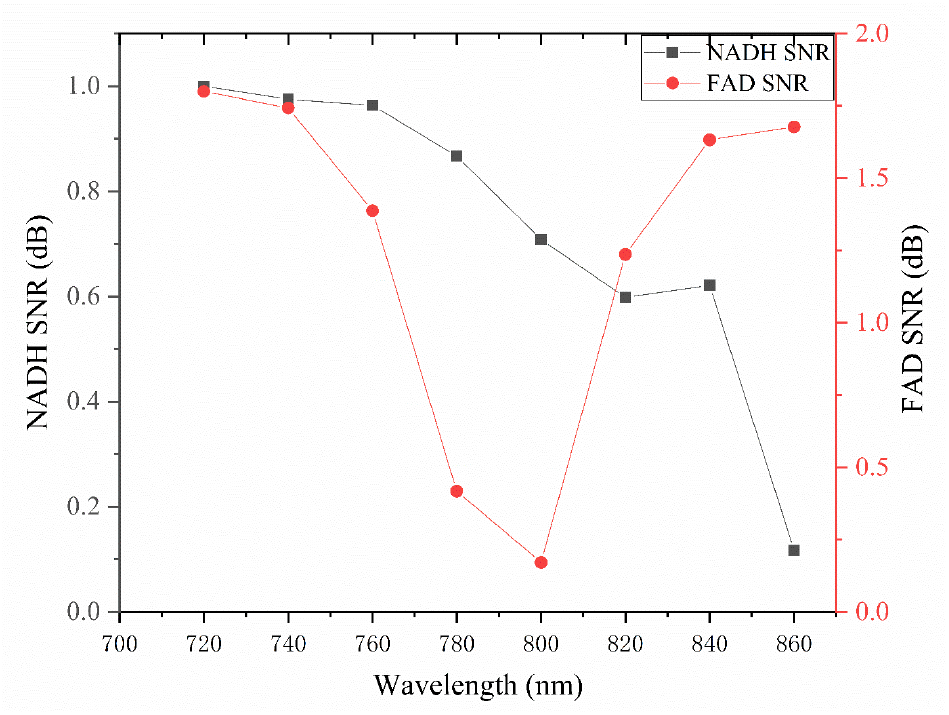
Performance about image quality with parameters of signal to noise ratio(SNR) in different wavelength from 720nm to 860nm. The unit is dB.

**Table S1.**
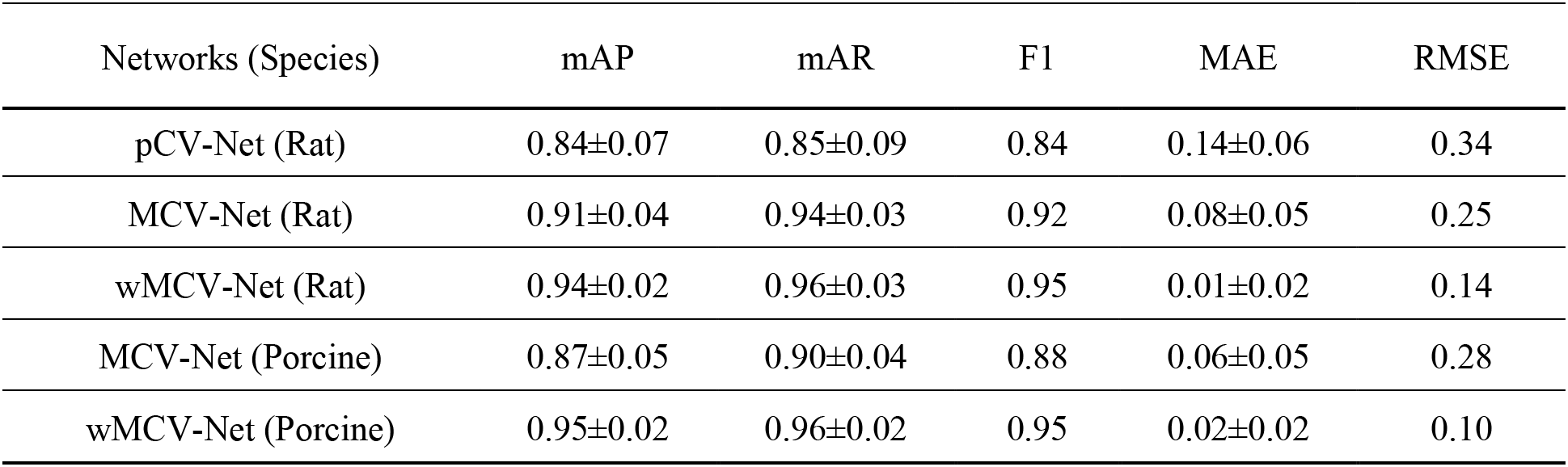
Table of result comparation with different deep learning algorithms in different animals.

| Networks (Species) | mAP | mAR | F1 | MAE | RMSE |
| --- | --- | --- | --- | --- | --- |
| pCV-Net (Rat) | 0.84±0.07 | 0.85±0.09 | 0.84 | 0.14±0.06 | 0.34 |
| MCV-Net (Rat) | 0.91±0.04 | 0.94±0.03 | 0.92 | 0.08±0.05 | 0.25 |
| wMCV-Net (Rat) | 0.94±0.02 | 0.96±0.03 | 0.95 | 0.01±0.02 | 0.14 |
| MCV-Net (Porcine) | 0.87±0.05 | 0.90±0.04 | 0.88 | 0.06±0.05 | 0.28 |
| wMCV-Net (Porcine) | 0.95±0.02 | 0.96±0.02 | 0.95 | 0.02±0.02 | 0.10 |

**Table. S2.** The accuracy of identifying chondrocytes with its live or dead status in different channels.

| Networks (Species) | Error rate (live cells) | Error rate (dead cells) | Error rate (total cells) |
| --- | --- | --- | --- |
| wMCV-Net (RG Porcine) | 0.34±0.38 | 0.53±0.37 | 0.27±0.20 |
| MCV-Net (RGB Porcine) | 0.23±0.20 | 0.24±0.30 | 0.22±0.30 |
| wMCV-Net (RGB Porcine) | 0.05±0.03 | 0.06±0.04 | 0.05±0.04 |

